# Enhanced Golic+: Gene targeting with 100% recovery in *Drosophila* male germ cells

**DOI:** 10.1101/681874

**Authors:** Hui-Min Chen, Xiaohao Yao, Qingzhong Ren, Chuan-Chie Chang, Ling-Yu Liu, Tzumin Lee

**Affiliations:** Howard Hughes Medical Institute, Janelia Research Campus, 19700 Helix Drive, Ashburn, VA 20147, USA

**Keywords:** *Drosophila*, Gene targeting, Homologous recombination, CRISPR, Male germline, Lethality-based selection

## Abstract

The efficiency of gene targeting can vary drastically. Even utilizing CRISPR/Cas9 does not ensure rapid, successful gene targeting. In *Drosophila*, we previously established Golic+ to augment gene-targeting productivity simply through fly pushing. This transgene-based system elicits gene targeting in germ cells. It further implements lethality selection to enrich for progeny with accurate gene targeting. However, limitations still remain. Here we deliver an improved Golic+ technique that we name Enhanced Golic+ (E-Golic+). E-Golic+ incorporates genetic modifications to eliminate false positives while simultaneously boosting efficiency. Strikingly, male germ cells are exceptionally susceptible to gene targeting using E-Golic+. With male founders, we easily achieve 100% recovery of correct gene targeting. Given the superior efficiency/specificity and relatively effortless scalability, E-Golic+ promises to triumph in any challenging and otherwise unattainable gene targeting projects in *Drosophila*.

## INTRODUCTION

The prokaryotic immune system, CRISPR/Cas9, has been successfully adopted for genome editing in diverse species (Komor et al. 2017). An engineered, widely used CRISPR/Cas9 system consists of two components: a single-molecule guide RNA (gRNA) and the Cas9 DNA endonuclease (Jinek et al. 2012; Hwang et al. 2013). The gRNA binds with Cas9 and directs Cas9 to produce double-strand DNA breaks in specific DNA sequences determined by base pairing between the gRNA and a 20bp DNA target. The only additional requirement in the DNA is the presence of protospacer adjacent motif (PAM, canonically NGG) immediately following the 20bp target sequence. One can therefore utilize CRISPR/Cas9 to target almost any genomic region with extremely high selectivity. The resultant DNA breaks are often repaired by non-homologous end joining (Lieber 2010), leading to deletions or (less frequently) insertions until the target sequence is lost. Notably, the likely indel profiles can be largely predicted based on local DNA sequences (Allen et al. 2018). The simplicity, robustness and predictability of Cas9-induced indels have made CRISPR as the most favored strategy for targeted gene disruption. Further, one can try to edit the genome around the Cas9 cut site via homology directed repair (HDR) of DNA breaks (San Filippo et al. 2008). With HDR, one can replace endogenous sequences with some designer sequences by supplying an exogenous template carrying the desired DNA sequence flanked by homology arms. Such tailored genome modifications are versatile but can be difficult if not impossible to achieve even with the CRISPR technology (Horlbeck et al. 2016; Isaac et al. 2016).

Gene targeting is context-dependent and offers little flexibility in the design. Some gene-targeting experiments are intrinsically more challenging than others. For instance, certain manipulations strive for deletion of a sizable defined DNA fragment or insertion of a long DNA sequence at a specific nucleotide position. This can be extremely challenging if suitable gRNA sites are not available. Moreover, it can be difficult to obtain and insert long homology arms into an already lengthy donor DNA. In addition, the engineered gene products (made through correct gene targeting) may unexpectedly compromise organism viability even in heterozygous conditions. To recover rare gene-targeting events in those challenging cases requires (1) generation of numerous offspring, each with independent trials, and (2) enrichment of offspring with correct gene targeting (especially those with decreased viability) by selection against ‘unperturbed’ progeny.

Golic+ is a transgene-based gene targeting system designed to achieve the above two objectives (Chen et al. 2015). First, it employs a *bam* promoter to confine gene targeting to germ cells rather than germline stem cells (Chen and McKearin 2003; Lehmann 2012). This theoretically guarantees independent gene targeting events in individual offspring. Second, it carries two conditional toxic genes: one to eliminate offspring that did not incorporate the donor DNA and the other to select against the incorporation of donor DNA in off-target sites. These lethality-based selections should therefore allow only offspring with correct gene targeting to survive into adults. We envisioned that a low probability gene-targeting event would occur eventually, and that assuming no recovery of false positives in Golic+, patience and simple fly pushing would be all that is needed to ensure success. The induction of gene targeting in germ cells further eliminates the need for single-founder crosses, a practice to avoid recovery of clonally identical lines. The amount of fly pushing can therefore be greatly reduced. Thus, for complex editing of genes in their native environment, Golic+ is particularly affordable compared to embryo injections.

Nonetheless, since its debut in 2015, the original Golic+ failed to succeed at all gene-targeting experiments. We suspended several trials due to the inability to recover correct gene targeting events after proving many candidates as false positives. In this study, we deliver an enhanced Golic+ (E-Golic+) with (1) much more stringent lethality selections plus (2) superior gene targeting efficiency. Strikingly, the E-Golic+ acts much more potently in male than female germ cells. From male founders, we easily achieve a 100% success-rate with previously failed gene-targeting experiments. Only in the most challenging case did we detect any false positives. In this case, offspring with off-target integration were outnumbered two-fold by offspring with correct gene targeting. Therefore, for extremely intractable or large-scale gene targeting experiments, one can perform group crosses to drastically reduce the labor required for making numerous independent trials with minimal false-positive contamination. In conclusion, E-Golic+ guarantees successful gene targeting in *Drosophila*.

## RESULTS

### Enhanced Golic+ reduces false positives while boosting efficiency

With Golic+, we can readily expand fly crosses to increase independent gene-targeting trials. However, despite lethality-based selections, most gene targeting trials yielded a significant number of false positives; and some Golic+ crosses produced very few survivors in total. We therefore re-examined the Golic+ design for potential shortcomings. In Golic+, a minimal *bam* promoter, bamP(198) co-expresses Cas9, FLP, and I-SceI in female germ cells. Cas9 directed by gRNA makes specific DNA cuts in the target gene. FLP mediates formation of the circular donor DNA from a pre-integrated FRT cassette, and I-SceI subsequently linearizes the donor. Golic+ further employs three LexA-dependent transgenes, including one repressible and one non-repressible toxic gene as well as a repressor gene, for lethality-based progeny selection. The repressible toxic gene exists in two parts separated by an FRT cassette that contains 5’ homology arm, the repressor gene, 3’ homology arm, and the non-repressible toxic gene in sequence. Excision of the FRT cassette would automatically reconstitute the repressible toxic gene at the original integration site of the donor DNA. We can then render organism survival contingent upon re-integration of the repressor gene. However, the liberated donor DNA carries the repressor gene as well as a non-repressible toxic gene. Given only the repressor gene flanked by homology arms, HDR-mediated gene targeting would naturally segregate the repressor gene from the non-repressible toxic gene and selectively place the repressor gene back to the genome. Thus, Golic+ permits enrichment of correct gene targeting.

Given the dependence of all key enzymes on the *bam* promoter, we first wondered if the strength of bamP(198) is a key limiting factor in the performance of Golic+. We addressed this issue by trying bamP(898), a longer and presumably stronger *bam* promoter (Chen and McKearin 2003). Notably, co-induction of Cas9, FLP, and I-SceI by bamP(898) yielded many more survivors including false positives at even higher ratios (Supplemental Table). The predominance of false positives overshadowed the evidently more potent bamP(898). To improve the efficiency of Golic+ we need to further identify and eliminate the source(s) of false positives.

We detected two categories of false positives. The first group consisted of escapers, those without donor DNA incorporation. Errors in the donor DNA liberation step resulted in defective reconstitution of the repressible toxicity gene (Fig. 1A). Without a functional repressible toxic module, organism viability was no longer coupled to genome incorporation of the donor DNA. To eliminate these escapers, we need to ensure presence of an intact, repressible toxic gene ideally at the same homologous site as the pre-integrated donor DNA. We met this requirement by making and placing the 3xP3-RFP-marked *3XLexAop2-riTS-Rac*^*V12*^ transgene at the same *attP* sites used for holding donor DNAs (Fig. 1B). This guarantees that all 3xP3-RFP-marked survivors carry an intact repressible toxic gene. Organism survival would therefore depend on relocation of the repressor-marked donor DNA onto a different (hopefully the desired) chromosome.

**Figure 1.**
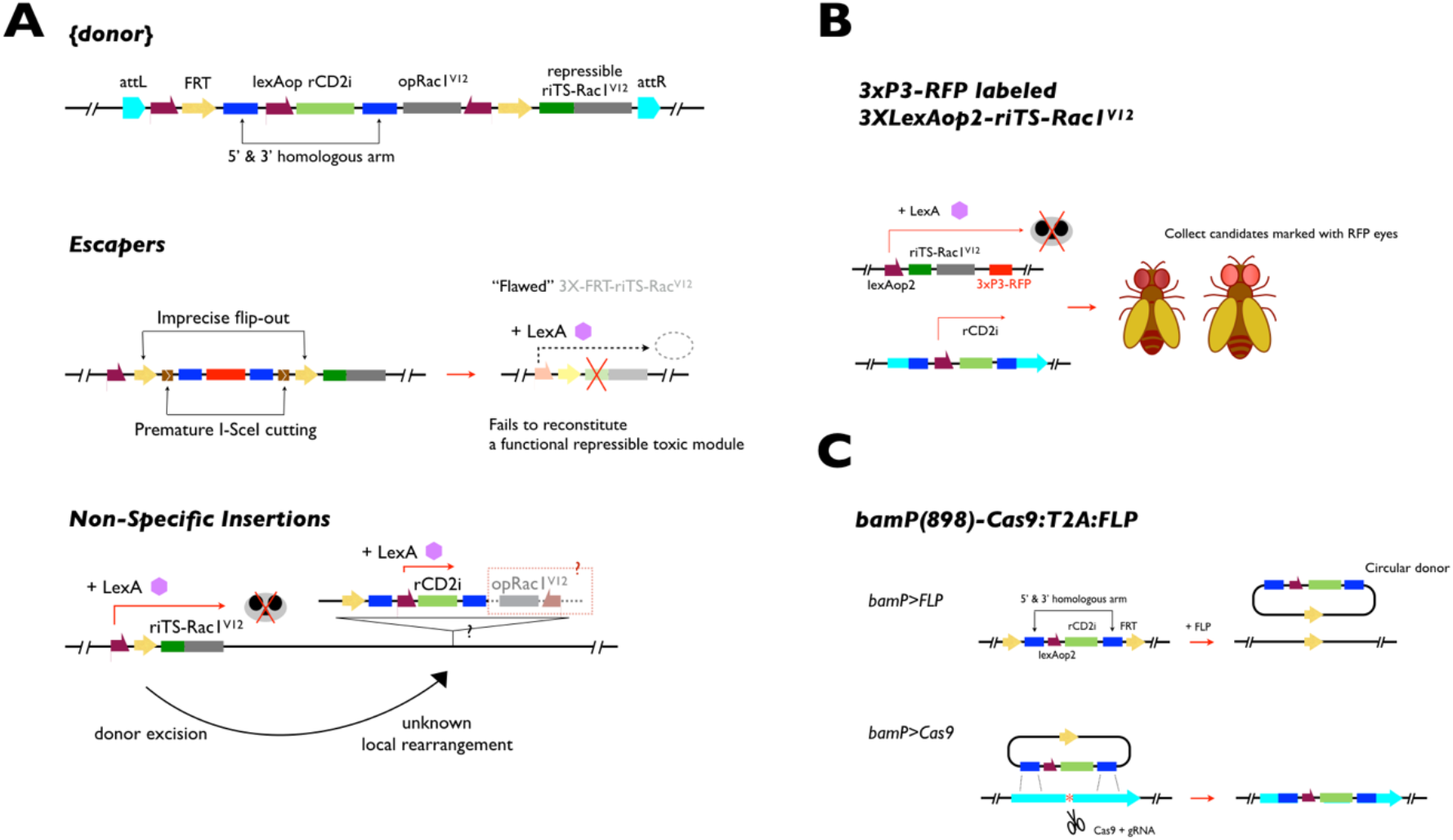
Improvements of E-Golic+ to remove two types of false positives, escapers and non-specific insertions. (A) In Golic+, {donor} was designed with two built-in toxicity modules and integrated in pre-characterized *attP* sites for efficient donor DNA release. We detected two false positive scenarios that produced progeny evading lethality selection. For escapers, they originated from failures in the reconstitution of a toxic module at the {donor} residual site, due to either imprecise flip-out or destructive premature I-SceI cutting. Therefore, they eclosed without ever being challenged by the lethality selection. For non-specific insertions, they retained the rCD2i suppressor and primarily relocated into the same chromosome. Yet they lost the ancillary non-repressible toxic module over this process, and survived the lethality selection without going through HDR. (B) We created 3xP3-RFP labeled *3XLexAop2-riTS-Rac1*^*V12*^ transgenic lines, and purposefully only collected surviving candidates marked with red fluorescent eyes. Hence, we effectively screened for candidates carrying the rCD2i suppressor, and avoid escapers completely. (C) Using *bamP(898)-Cas9:T2A:FLP*, we induced HDR in germ cells with CRISPR and circular donor DNA, hence directly relieved ourselves from the occurrence of non-specific insertions originating from linear donor DNA.

The second group of false positives resulted from non-specific insertions of the donor DNA. Per Golic+ design, HDR at the correct target site should segregate the repressor and the non-repressible toxic gene, as they are separated by one of the paired homology arms. By contrast, organisms with non-specific insertions should retain the non-repressible toxic gene and fail to survive upon selection with some broad LexA driver. However, the non-specific insertions we recovered had somehow selectively lost the non-repressible toxic gene (Fig. 1A). While we do not know how this occurred, we may be able to better preserve the integrity of the liberated donor DNA by keeping it in a circular form. Further, linear DNA can promote non-specific insertion and circular DNA is competent as a template for HDR (Beumer et al. 2008). To this end, we made *bamP(898)-Cas9:T2A:FLP* that drives only Cas9 and FLP, thus excluding I-SceI (Fig. 1C). We refer to Golic+ with *bamP(898)-Cas9:T2A:FLP* plus 3xP3-RFP-marked *3XLexAop2-riTS-Rac*^*V12*^ as Enhanced Golic+. Please see Table 1 for transgenes required for implementing enhanced Golic+ and Figure 2 for representative targeting schemes.

**Table 1.**
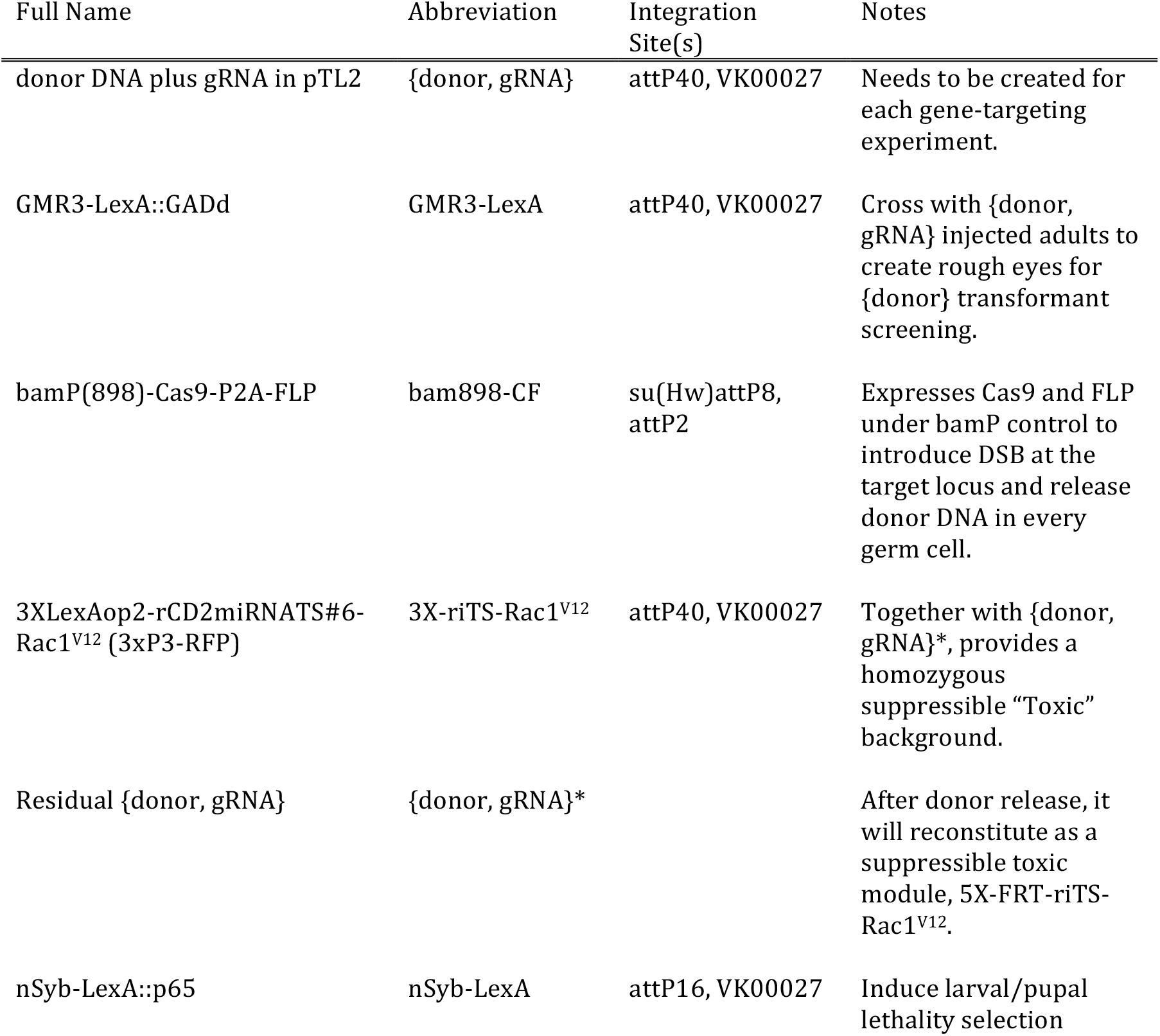
List of transgenic lines required for implementing Enhanced Golic+.

**Figure 2.**
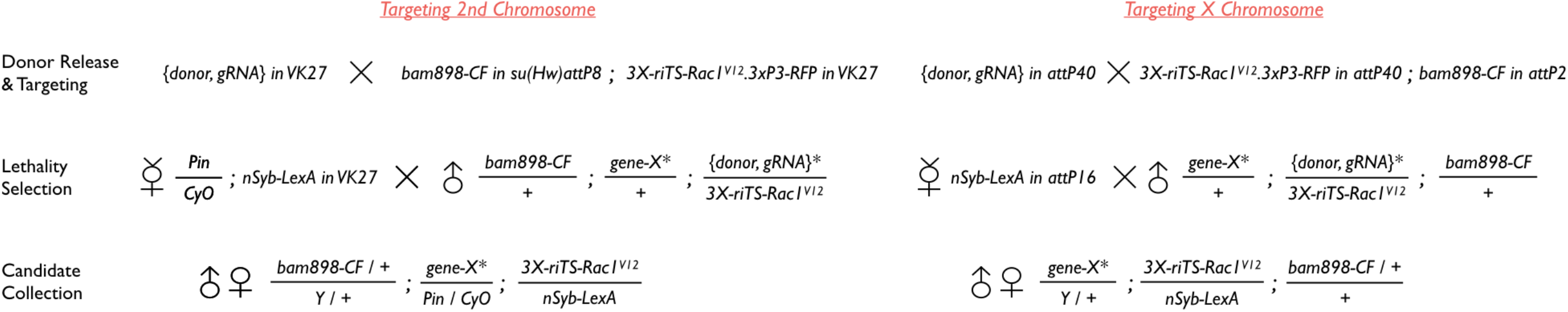
Targeting schemes for a second or an X chromosome gene. Like Golic+, E-Golic+ involves two crosses and three steps. In the first cross, we create founders that have active CRISPR reactions with circular donor for HDR in their germ cells. Then, founders are mated with *nSyb-LexA* so that each progeny will experience lethality selection, and most, if not all, of the 3xP3-RFP marked surviving candidates inherit gene targeting events marked with rCD2i suppressor.

We performed a direct comparison of Golic+ with E-Golic+ to see if we could eliminate false positives and increase efficiency. Using enhanced Golic+, we effectively eliminated all false positives observed in three previously failed Golic+ experiments (Fig. 3). We were further able to recover multiple correct gene-targeting events in one of the three genes we tested. These results substantiate the success in eliminating false positives with the newly introduced transgenes plus use of circular templates instead. However, two of the three repeated trials remained unsuccessful, demanding larger scales of fly pushing or higher gene targeting efficiencies.

**Figure 3.**
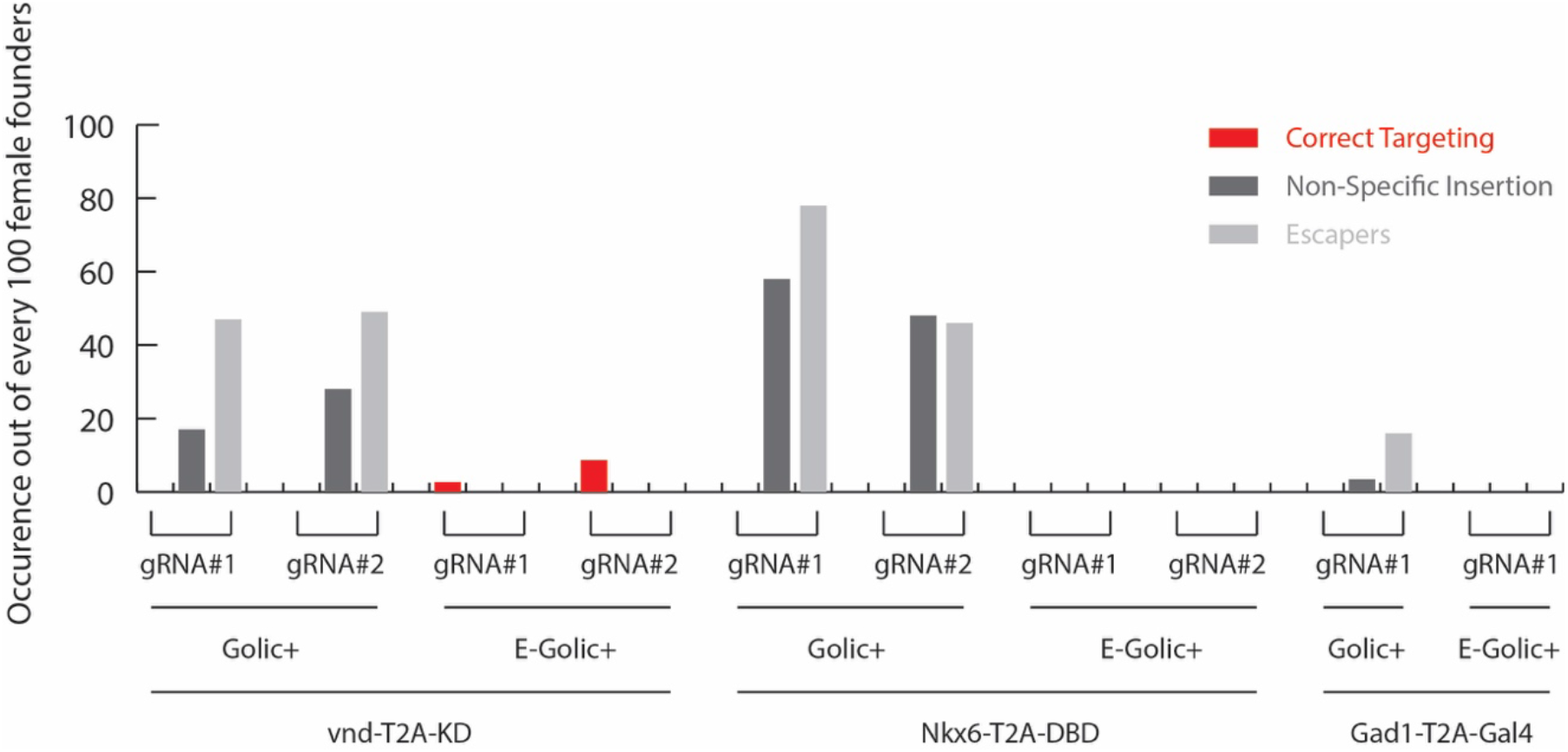
E-Golic+ effectively eliminates the occurrence of false positives. To evaluate the new transgenes introduced in E-Golic+, we performed gene targeting with five different donors using either Golic+ or E-Golic+. Occurrence of three different types of gene targeting candidates (correct targeting, non-specific insertion, and escapers) out of every 100 female founders are numbers interpolated or extrapolated from data in Table 2.

### Males make superior founders

One laborious step of performing E-Golic+ is the collection of copious virgin females to be founders. Conversely, using males as founders would significantly reduce the load of fly pushing when many founders are needed to obtain rare gene targeting events. Males should be able to be used as founders as *bam* shows comparable restricted expression in both male and female gonads (Fuller and Spradling 2007). Hence, use of *bamP898* in E-Golic+ should also effectively confine gene targeting to male germ cells. We therefore repeated all three gene-targeting experiments with E-Golic+ in male founders.

**Table 2.**
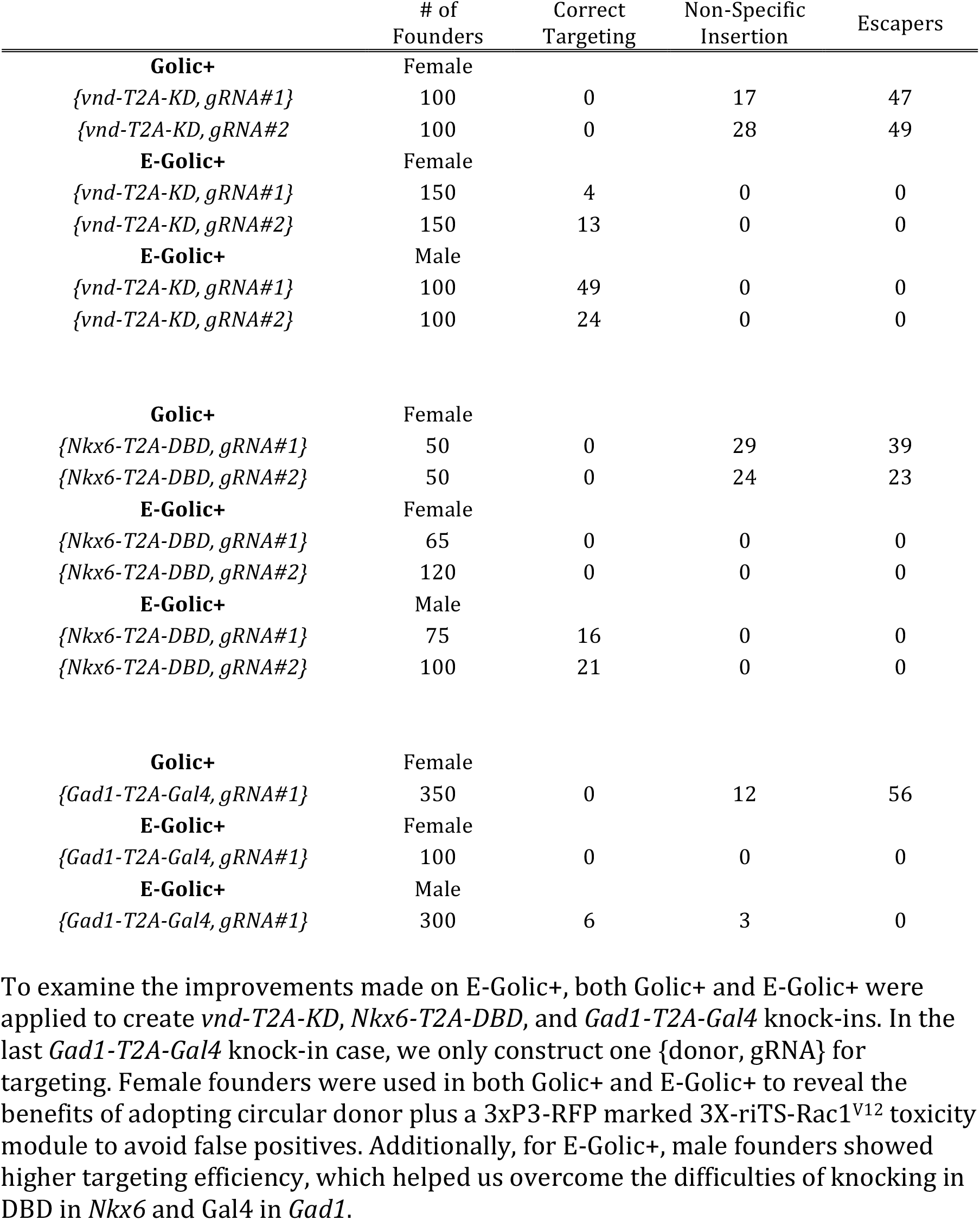
Comparison of Golic+ and Enhanced Golic+.

Using male founders, we readily recovered numerous correct gene-targeting offspring from each of the three gene-targeting trials (Table 2). None of these trials were successful with Golic+, and only one was successful with E-Golic+ using female founders. To make *vnd:T2A:KD*, we recovered 73 offspring with *vnd:T2A:KD* from a total of 200 male founders, as opposed to only 17 from a total of 300 female founders. In the engineering of *Nkx6:T2A:DBD*, we utilized two gRNA choices and obtained 37 offspring with *Nkx6:T2A:DBD* from a total of 175 male founders, but recovered none from a total of 185 female founders. In the third case, we aimed to insert Gal4 into *Gad1*, which encodes an enzyme characteristic of GABAergic neurons, to make *Gad1:T2A:Gal4*. Expressing GAL4 continuously in all GABAergic neurons could be harmful. In fact, an earlier study has reported challenges in maintaining an analogous fly stock generated through recombinase-mediated cassette exchange (Diao et al. 2015). Given the known challenges, we screened through progeny from 300 male founders and recovered six offspring with *Gad1:T2A:Gal4*. We validated the lines carrying *Gad1:T2A:Gal4* by genomic PCR, and further corroborated their Gal4 expression patterns highlighting GABAergic neurons in adult brains co-stained with anti-GABA antibody (Fig. 4). As expected, we found that *Gad1:T2A:Gal4* labeled several prominent groups of GABAergic neurons reported previously (Okada et al. 2009). We observed prominent labeling of R neurons that innervate the ellipsoid body (Fig. 4A1), neurons dorsal, ventral, and lateral to the antennal lobe neuropil (Fig. 4A2), neurons on the surface of Medulla (Fig. 4A3), and neurons at the interface between medulla and lobula plate in the posterior brain (Fig. 4A4). In addition to six correct gene-targeting lines carrying *Gad1:T2A:Gal4*, we found three false positives with non-specific insertions. It is unclear how a portion of circular DNA that contains the repressor became integrated into the genome in these flies. However, the drastic suppression of non-specific insertion in E-Golic+ is evident, indicating the importance of using extra-chromosomal circular DNAs as templates for HDR.

In sum, the enhanced Golic+ is particularly powerful in the male germline. The lethality-based selections against false positives remain highly stringent. Moreover, the efficiency of gene targeting is greatly enhanced such that we could readily recuse all previously failed Golic+ experiments with E-Golic+.

**Figure. 4.**
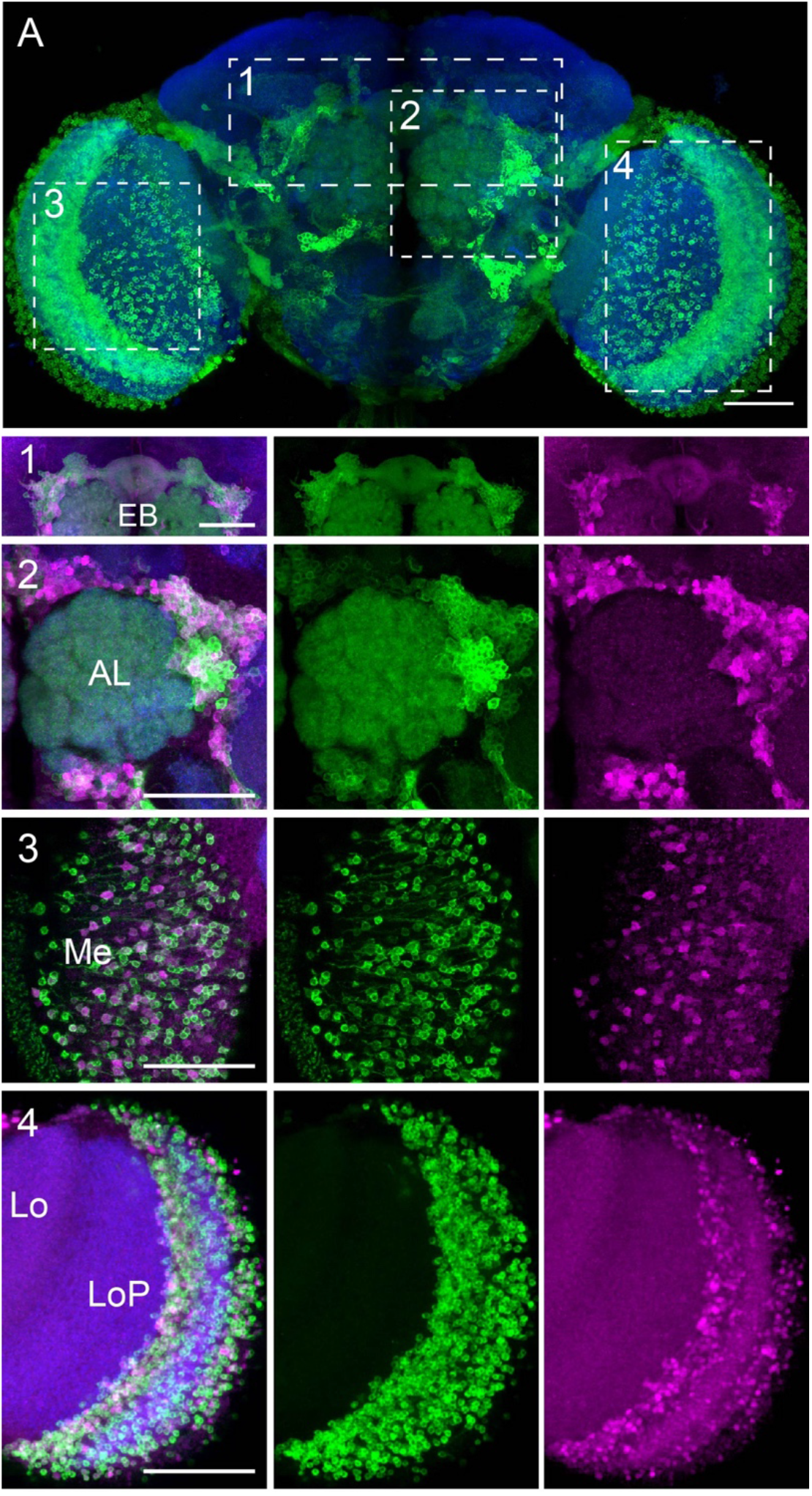
The expression pattern of *Gad1:T2A:Gal4* in *Drosophila* central nervous system. (A) Composite confocal images of an adult fly brain with *Gad1:T2A:Gal4* driving a neuronal membrane marker (10XUAS-mCD8-GFP, green). The brain was counterstained with an anti-Bruchpilot protein antibody which specifically label presynaptic active zones (blue). Partial projections of the boxed regions in (A) were shown separately below together with anti-GABA staining (Magenta). 1: The Ellipsoid Body (EB) region; 2: The Antennal Lobe (AL) region; 3: The Medulla (Me) surface; 4: The interface between Medulla and Lobula Plate (LoP). Lo: Lobula. Scale bar, 50um in all panels.

## DISCUSSION

Homology-dependent gene targeting allows designer genome editing, but suffers from unpredictable success even with modern CRISPR/Cas9 technology. By tackling previously failed gene-targeting trials, we show that the E-Golic+ reaches a 100% success in gene targeting experiments. There are two levels of enhancement. First, E-Golic+ offers absolute lethality selection to expedite the recovery of correct gene targeting. Second, E-Golic+ achieves an exceptionally high efficiency of gene targeting in male germline. E-Golic+ is probably the most sophisticated gene targeting system to date. It amplifies the power of *Drosophila* genetics and exemplifies how to access and modify the genomes of higher organisms.

Past studies have shown higher levels of gene targeting in female germline (Rong and Golic 2000), but more efficient targeted mutagenesis in male germline (Bibikova et al. 2002). Gene targeting depends on homologous recombination, while gene disruption occurs through non-homologous repair. Such mechanistic distinctions have promoted the idea that the lack of meiotic homologous recombination in *Drosophila* male germline may underlie the previously published gender differences in gene targeting versus gene disruption. However, our data suggest that male germ cells are much more susceptible than female germ cells to Cas9-mediated genome editing via HDR. These differences might result from repairs of double-strand DNA breaks in germline stem cells versus germ cells. Interestingly, a recent paper reported that CRISPR-induced DSBs can be repaired through recombination across homologous chromosomes in germline stem cells (Brunner et al. 2019). We further speculate that the homologous chromosomes in male germ cells might not be intimately paired for recombination and thus individually more susceptible to repairs by donor DNA. Nonetheless, we establish male germ cells as the top choice for germline genome editing by E-Golic+ in *Drosophila*.

In our efforts to eliminate false positives, we confirmed that one could effectively prevent off-target integration of the liberated donor DNA by keeping it in the intact circular form. Once linearized, the donor DNA becomes prone to non-specific insertion. Notably, the rate of non-specific insertion for linearized donor DNA varies depending on the donor. Seemingly, there is an inverse correlation between the non-specific insertion rate and the success rate of gene targeting. By contrast, it appears that the off-target integration of circular donor DNAs remains persistently suppressed regardless of actual gene-targeting efficiency. These phenomena implicate differential fates for linear versus circular extra-chromosomal DNA, further elucidation of which may help improve future gene targeting.

In sum, E-Golic+ in male germ cells has triumphed in previously failed gene-targeting experiments with a 100% success rate. Impressively, almost 100% of recovered candidates carried the desired genome modifications at the correct site. Moreover, to achieve really intractable gene targeting, one can readily continue the attempts by simple fly pushing. Given its unparalleled efficiency, specificity, and scalability, enhanced Golic+ in male germ cells promises to enable further sophisticated genome editing in *Drosophila* and beyond.

## MATERIAL AND METHODS

### Fly strains

Here are the fly strains used in this study: (1) *bamP(198)-Cas9:2A:FLP:2A:I-SceI* in *su(Hw)attP8* and *attP2*; (2) *bamP(898)-Cas9:2A:FLP:2A:I-SceI* in *su(Hw)attP8* and *attP2*; (3) *GMR3-LexA::GADd* in *attP40* and *VK00027*; (4) *nSyb-LexA::p65* in *attP16* and *VK00027*; (5) *bamP(898)-Cas9:2A:FLP* in *su(Hw)attP8* and *attP2*; (6) *3X-riTS-Rac1V12(3xP3-RFP)* in *attP40* and *VK00027.*

### Molecular biology

To create *vnd-T2A-KD*, *Nkx6-T2A-DBD*, and *Gad1-T2A-Gal4* knock-ins, 5’ and 3’ homology arms of approximately 1.5 kb in length and right before or after the *vnd*, *Nkx6*, and *Gad1* stop codons were amplified from genomic DNA and cloned into pTL2. Homology arms were further mutated to avoid CRISPR cutting on the donor. The following CRISPR target sites were chosen: vnd_gRNA#1: GCATGGCCGTGCAGTAGACC; vnd_gRNA#2: GTTCCTCACCAGAACTGGAA; Nkx6_gRNA#1: GAAATTAAGTCTTCAGAAGA; Nkx6_gRNA#2: GCCATTTGGTGCGACGATTC; Gad1_gRNA#1: GCTACCAGCCCGACGATCGC. T2A-KD and T2A-DBD were introduced by cloning KD and DBD from pJFRC161-20XUAS-IVS-KD::PEST (Nern et al. 2011) and pBPZpGAL4DBDUw (Pfeiffer et al. 2010). The full *bam* promoter (−898) (Chen and McKearin 2003) was ordered from gBlocks, IDT to create bamP(898)-Cas9:2A:FLP:2A:I-SceI. Afterwards, coding sequence of Cas9:2A:FLP:2A:I-SceI was replaced by a PCR amplification of only the Cas9:2A:FLP portion to generate bamP(898)-Cas9:2A:FLP.

### Fly genetics

{vnd-T2A-KD, gRNA#1}, {vnd-T2A-KD, gRNA#2}, {Nkx6-T2A-DBD, gRNA#1}, {Nkx6-T2A-DBD, gRNA#2}, and {Gad1-T2A-Gal4} were all integrated in attP40 to target *vnd* on the X chromosome, *Nkx6* and *Gad1* on the third chromosome. Transgenic *{donor, gRNA}* donors were mated with flies carrying *bam898-CF* and *3X-riTS-Rac1V12(3xP3-RFP)* to create E-Golic+ founders. These founders were then crossed to *nSyb-LexA* flies for lethality selection. Finally, for E-Golic+, only Surviving candidates labeled with 3xP3-RFP were subjected to chromosomal mapping and genomic PCR confirmation.

### Immunostaining and fluorescence microscopy

We dissected adult fly brains in ice-cold phosphate-buffered saline (PBS) and immediately transferred them into 4% paraformaldehyde for fixation at room temperature. After 30min fixation and three washes in PBS plus 0.5% Triton-X-100 (PBT), we added blocking solution (PBT with 4% Normal Goat Serum) and blocked the brains for 1 hour. Next, we transferred the brains into blocking solution containing primary antibodies and incubated at 4 °C overnight. After three 30-min wash in PBT, we added secondary antibodies in blocking solution and incubated for two days. Finally, after washing three additional times in PBT, we transferred the brains into PBS and mounted in SlowFade Gold Reagent on charged slides (Fisherbrand, 12-550-15).

Primary antibodies include: Chicken anti-GFP (1:1000; Life Technologies, A10262), Rabbit anti-GABA (1:25; Millipore Sigma, A2052), and mouse anti-nc82 (1:40; Developmental Studies Hydridoma Bank or DSHB). Secondary antibodies include: Alexa-488-conjugated goat antibody to chicken (1:500; ThermoFisher Scientific, A-11039), Cy3-conjugated goat antibody to Rabbit (1:200; Jackson ImmunoResearch, #111-165-144), and Cy5-conjugated goat antibody to mouse (1: 200; Jackson ImmunoResearch, #115-605-146).

We acquired image stacks of whole-mount fly brains using a Zeiss LSM 710 confocal microscope. The images were taken at 1um intervals at 1024×1024 pixel resolution using a 40X C-Apochromat water objective (NA=1.2). The images were further processed with Fiji and Adobe Photoshop.

## Acknowledgements

We thank Dr. Fillip Port and Dr. Kate Koles for helpful discussion. We thank Dr. Rosa Linda Miyares for critical editing of the manuscript. We thank Janelia Fly Core for technical support. We thank Crystal Di Pietro and Kathryn Miller for administrative support. This work was supported by Howard Hughes Medical Institute.

## Author contributions

H.-M.C. and T.L. conceived the project. H.-M.C. performed the experiments. X.Y. and C.-C.C. generated E-Golic+ constructs. Q.R. analyzed the Gad1-T2A-Gal4 expression patterns. L.-Y.L. assisted in E-Golic+ screening. H.-M.C. and T.L. wrote the manuscript. T.L. supervised the project.

---

**Supplemental Table.** Golic+, comparing *bam198-CFI* and *bam898-CFI* with *{msh-T2A-Gal4 KI, gRNA}*.

|  | # of Female Founders | Correct Targeting | Non-Specific Insertion | Escapers |
| --- | --- | --- | --- | --- |
| *bam198-CFI* | 95 | 47 | 8 | 15 |
| *bam898-CFI* | 30 | 37 | 53 | 5 |

Using a common donor *{msh-T2A-Gal4 KI, gRNA}*, we compare the performance of *bam198-CFI* and *bam898-CFI* in terms of the production of correct targeting, non-specific insertion, and escaper candidates.

## References

Allen F, Crepaldi L, Alsinet C, Strong AJ, Kleshchevnikov V, De Angeli P, Palenikova P, Khodak A, Kiselev V, Kosicki M. et al. 2018. Predicting the mutations generated by repair of Cas9-induced double-strand breaks. Nat Biotechnol.

Beumer KJ, Trautman JK, Bozas A, Liu J-L, Rutter J, Gall JG, Carroll D. 2008. Efficient gene targeting in Drosophila by direct embryo injection with zinc-finger nucleases. Proceedings of the National Academy of Sciences 105: 19821–19826.

Bibikova M, Golic M, Golic KG, Carroll D. 2002. Targeted chromosomal cleavage and mutagenesis in Drosophila using zinc-finger nucleases. Genetics 161: 1169–1175.

Brunner E, Yagi R, Debrunner M, Beck-Schneider D, Burger A, Escher E, Mosimann C, Hausmann G, Basler K. 2019. CRISPR-induced double-strand breaks trigger recombination between homologous chromosome arms. Life science alliance 2.

Chen D, McKearin DM. 2003. A discrete transcriptional silencer in the bam gene determines asymmetric division of the Drosophila germline stem cell. Development 130: 1159–1170.

Chen HM, Huang Y, Pfeiffer BD, Yao X, Lee T. 2015. An enhanced gene targeting toolkit for Drosophila: Golic+. Genetics 199: 683–694.

Diao F, Ironfield H, Luan H, Diao F, Shropshire WC, Ewer J, Marr E, Potter CJ, Landgraf M, White BH. 2015. Plug-and-play genetic access to drosophila cell types using exchangeable exon cassettes. Cell reports 10: 1410–1421.

Fuller MT, Spradling AC. 2007. Male and female Drosophila germline stem cells: two versions of immortality. Science 316: 402–404.

Horlbeck MA, Witkowsky LB, Guglielmi B, Replogle JM, Gilbert LA, Villalta JE, Torigoe SE, Tjian R, Weissman JS. 2016. Nucleosomes impede Cas9 access to DNA in vivo and in vitro. eLife 5.

Hwang WY, Fu Y, Reyon D, Maeder ML, Tsai SQ, Sander JD, Peterson RT, Yeh JJ, Joung JK. 2013. Efficient genome editing in zebrafish using a CRISPR-Cas system. Nature biotechnology 31: 227–229.

Isaac RS, Jiang F, Doudna JA, Lim WA, Narlikar GJ, Almeida R. 2016. Nucleosome breathing and remodeling constrain CRISPR-Cas9 function. eLife 5.

Jinek M, Chylinski K, Fonfara I, Hauer M, Doudna JA, Charpentier E. 2012. A programmable dual-RNA–guided DNA endonuclease in adaptive bacterial immunity. Science 337: 816–821.

Komor AC, Badran AH, Liu DR. 2017. CRISPR-Based Technologies for the Manipulation of Eukaryotic Genomes. Cell 169: 559.

Lehmann R. 2012. Germline stem cells: origin and destiny. Cell stem cell 10: 729–739.

Lieber MR. 2010. The mechanism of double-strand DNA break repair by the nonhomologous DNA end-joining pathway. Annual review of biochemistry 79: 181–211.

Nern A, Pfeiffer BD, Svoboda K, Rubin GM. 2011. Multiple new site-specific recombinases for use in manipulating animal genomes. Proceedings of the National Academy of Sciences of the United States of America 108: 14198–14203.

Okada R, Awasaki T, Ito K. 2009. Gamma-aminobutyric acid (GABA)-mediated neural connections in the Drosophila antennal lobe. The Journal of comparative neurology 514: 74–91.

Pfeiffer BD, Ngo T-TB, Hibbard KL, Murphy C, Jenett A, Truman JW, Rubin GM. 2010. Refinement of tools for targeted gene expression in Drosophila. Genetics 186: 735–755.

Rong YS, Golic KG. 2000. Gene targeting by homologous recombination in Drosophila. Science 288: 2013–2018.

San Filippo J, Sung P, Klein H. 2008. Mechanism of eukaryotic homologous recombination. Annual review of biochemistry 77: 229–257.

